# Metabolic Modulation Predicts Heart Failure Tests Performance

**DOI:** 10.1101/555417

**Authors:** Daniel Contaifer, Leo F. Buckley, George Wohlford, Naren G Kumar, Joshua M. Morriss, Asanga D. Ranasinghe, Salvatore Carbone, Justin M Canada, Cory Trankle, Antonio Abbate, Benjamin W. Van Tassell, Dayanjan S Wijesinghe

## Abstract

Cardiopulmonary testing (CPET) and biomarkers such as NT-proBNP, Galectin-3, and C-reactive protein (CRP) have shown great promise for characterizing the heart failure (HF) phenotypes. However, the underlying metabolic changes that cause, predict and accompany changes in these parameters are not well known. In depth knowledge of these metabolic changes has the ability to provide new insights towards the clinical management of HF as well as providing better biomarkers for the assessment of disease progression and measurement of success from clinical interventions. To address this knowledge gap, we undertook a deep metabolomic and lipidomic phenotyping of a cohort of HF patients and utilized Multiple Regression Analysis (MRA) to identify associations between the patient’s plasma metabolome and the lipidome to parameters of CPET and the above mentioned HF biomarkers (HFBio). A total of 49 subjects were submitted to CPET and HFBio evaluation with deep metabolomic and lipidomic profiling of the plasma. Stepwise MRA and Standard Least Squares methods were used to determine models of best fit. Metabolic Pathway Analysis was utilized to detect what metabolites were over-represented and significantly enriched, and Metabolite Set Enrichment Analysis was used to explore preexisting biological knowledge from studies of human metabolism. The study cohort was predominantly males of African American decent, presenting a high frequency of diabetes, hyperlipidemia, and hypertension with a low mean left ventricular ejection fraction and a high left ventricular end-diastolic and end-systolic volume. VE/VCO2 and Peak VO2 (CV=0.07 and 0.08, respectively) showed the best metabolic predictive model for test performance, while CRP and NT-ProBNP (CV=0.31 and 0.29, respectively) were the least fitted models. Aminoacyl-tRNA and amino acid biosynthesis, amino acid metabolism, nitrogen metabolism, pantothenate and CoA biosynthesis, sphingolipid and glycerolipid metabolism, fatty acid biosynthesis, glutathione metabolism, and pentose phosphate pathway (PPP) were revealed to be the most relevant pathways. Principal related diseases were brain dysfunction and enzyme deficiencies associated with lactic acidosis. Considering the modulations in HF poor test performance, our results indicate an overall profile of oxidative stress, lactic acidosis, and metabolic syndrome coupled with mitochondria dysfunction. Overall, the insights resulting from this study’s MRA coincides with what has previously been discussed in parts in existing literature thereby confirming the validity of our findings while at the same time thoroughly characterizing the metabolic underpinning of CPET and HFBio results.

## 2. Introduction

The prevalence of heart failure (HF) has increased over time with the aging population. In people older than 20, the incidence of HF has increased from 5.7 million Americans between 2009 and 2012 to 6.5 million Americans by 2011. Despite aggressive measures to improve HF management, the five-year mortality rate of HF patients remains approximately 40%--comparable to many forms of cancer[1,2]. Investigations into diagnosis of HF has revealed promising cardiopulmonary tests and biomarkers that allow better disease management following diagnosis[3,4]. Combining patient’s metabolic profiles from HF compromised organs and tissues with HF tests has been demonstrated to provide physicians with an efficient source of clinical information used to both manage and diagnose patients[5]. However, the complex association of these HF tests with changes in the peripheral metabolism of compromised individuals is still under insufficient investigation and has failed to reveal the value of circulating metabolites as HF biomarkers[6].

Impaired cardiorespiratory fitness measured during cardiopulmomary exercise testing (CPET) is a hallmark manifestation of heart failure[7] and exercise training reduces all-cause mortality in patients with heart failure and reduced left ventricular ejection fraction (HFrEF)[8]. As such, cardiopulmonary exercise testing (CPET) is an evidence-supported assessment technique routinely used in the functional diagnosis of HF, prognostic evaluation of patients with chronic Heart Failure (CHF), and is also clinically relevant to supplement other clinical data in patients’ screening for heart transplantation[9,10]. Similarly, HF biomarkers (HFBio) such as NT-proBNP, Galectin-3, and C-reactive protein (CRP) have shown promising results as predictors of mortality[9,11]. However, a potential disadvantage of the CPET and HFBio use is that the evaluation of their response alone is insufficient to indicate risk of heart disease nor is it enough to diagnose a heart problem[12,13].

It is now known that cardiac and peripheral metabolic abnormalities may contribute to the pathogenesis of heart failure[14]. Studies of the metabolic profile of HF patients have indicated a rich metabolic modulation that can be identified and used as putative biomarkers[15–17]. However, little is known about the relationship of cardiorespiratory fitness and global metabolic profile in HFrEF. Efforts to correlate clinical and metabolic data are still necessary to fully integrate metabolomics as a translational medicine apparatus[18]. We hypothesized that deep metabolomic and lipidomic phenotyping would reveal novel metabolic and lipid mediators of cardiorespiratory fitness in patients with HFrEF. To test this hypothesis, we employed a Multiple Regression Analysis (MRA) study on the association of both CPET and HFBio with respects to the plasma lipidome and the metabolome of HF patients demonstrating that the metabolic modulation in HF patients depends on the tests’ performance. Although MRA cannot with certainty imply causality for the HF performance, this analysis intends to reveal the metabolic changes underlying the complex HF pathology to maximize the relevance of the HF tests and its potential for HF prognosis.

## 3. Materials and Methods

### 3.1) Patients

A post-hoc analysis was performed on patients who were enrolled in the REDHART study[19], which included patients with a left ventricular ejection fraction < 50% and high-sensitivity C-reactive protein (hsCRP) ≥2 mg/L and who were recently discharged after a hospitalization for HF. The present analysis includes plasma samples collected from 49 patients at baseline prior to randomization. The design and results of REDHART have been reported previously [19]. All patients underwent maximal aerobic exercise testing using a metabolic cart and a treadmill according to a conservative ramping treadmill protocol. Patients exercised to exhaustion (peak respiratory exchange ratio ≥1.00 and preferably ≥1.10) and those who were unable to complete the cardiopulmonary exercise test were excluded. The core CPET lab at University of Illinois at Chicago conducted all analyses of cardiopulmonary exercise test variables (peak oxygen consumption, oxygen uptake efficiency slope, exercise duration, minute ventilation-carbon dioxide production [VE/VCO2] as described previously[19].

### 3.2) Lipidomic and Metabolomic Data Acquisition

#### 3.2a) Lipid and metabolite extraction for LC-MS/MS analyses

Blood plasma lipids extraction was carried out using a biphasic solvent system of cold methanol, methyl tert-butyl ether (MTBE), and water with some modifications[20]. In detail, 225 μL of cold methanol containing a mixture of odd chain and deuterated lipid internal standards [lysoPE(17:1), lysoPC(17:0), PC(12:0/13:0), PE(17:0/17:0), PG(17:0/17:0), sphingosine (d17:1), d7 cholesterol, SM(17:0), C17 ceramide, d3 palmitic acid, MG(17:0/0:0/0:0), DG(18:1/2:0/0:0), DG(12:0/12:0/0:0), and d5 TG(17:0/17:1/17:0)] was added to a 20 μL blood plasma aliquot in a 1.5 mL polypropylene tube, and then vortexed. Next, 750 μL of cold MTBE was added, followed by vortexing and shaking with an orbital mixer. Phase separation was induced by adding 188 μL of MS-grade water. Upon vortexing (20s) the sample was centrifuged at 12,300 rpm for 2 min. The upper organic phase was collected in two 300 μL aliquots and evaporated with a vapor trap. Dried extracts were resuspended using 110 μL of a methanol/toluene (9:1, v/v) mixture containing CUDA (50 ng/ml; internal standard for quality control of injection) with support of vortexing (10 s), and centrifuged at 800 rpm for 5 min, followed by transferring 100 uL of the supernatant into autosampler vial with an insert. The lower polar layer was collected for metabolite analysis via HILIC LC-MS/MS method. An aliquot of 125 μL was evaporated to dryness in a SpeedVac and resuspended in acetonitrile.

#### 3.2b) Metabolomics:GC-MS metabolite extraction

30μl of plasma sample was added to a 1.0 mL of pre-chilled (−20°C) extraction solution composed of acetonitrile, isopropanol and water (3: 3: 2, v/v/v). Sample were vortexed and shaken for 5 min at 4°C using the Orbital Mixing Chilling/Heating Plate. Next, the mixture was centrifuged for 2min at 14,000 rcf. 450μL of the supernatant was dried with cold trap concentrator. The dried aliquot was then reconstituted with 450 μL acetonitrile:water (50:50) solution and centrifuged for 2 min at 14,000 rcf. The supernatant was transferred to a polypropylene tube and subjected to drying in a cold trap. The process of derivatization began with addition of 10 μL of 40 mg/mL Methoxyamine hydrochloride solution to each dried sample and standard. Samples were shaken at maximum speed at 30 ^o^C for 1.5 hours. Then, 91 μL of MSTFA + FAME mixture was added to each sample and standard and capped immediately. After shaking at maximum speed at 37 ^o^C, the content was transferred to glass vials with micro-serts inserted and cap immediately.

#### 3.2c) Lipids: LC-MC/MC conditions

Untargeted lipid analysis was obtained with Sciex TripleTOF 6600 coupled to Agilent 1290 LC. Lipids were separated on an Acquity UPLC CSH C18 column (100 × 2.1 mm; 1.7 μm) (Waters, Milford, MA, USA). The column was maintained at 65 °C and the flow-rate of 0.6 mL/min. The mobile phases consisted of (A) 60:40 (v/v) acetonitrile:water with 10 mM ammonium acetate and (B) 90:10 (v/v) isopropanol:acetonitrile with 10 mM ammonium acetate. The separation was conducted following a stepwise gradient: 0-2 min 15-30% (B), 2-2.5 min 48% (B), 2.5-11 min 82% (B), 11-11.5 min 99% (B), 11.5-12 min 99% (B), 12-12.1 15% (B), 12-14 min 15% (B). Negative and positive electrospray ionization (ESI) modes were applied with nitrogen serving as the desolvation gas and the collision gas. The parameters for detection of lipids were as follows: Curtain Gas: 35; CAD: High; Ion Spray Voltage: 4500 V; Source Temperature: 350°C; Gas 1: 60; Gas 2: 60; Declustering Potential: +/− 80V, and collision energies +/− 10.

#### 2.2d) Metabolites HILIC: LC-MS/MS conditions

Detection of water soluble plasma metabolites was carried out on Sciex TripleTOF 6600 coupled to Agilent 1290 LC. Analytes were separated on an Acquity UPLC BEH Amide Column, 130Å, 1.7 μm, 2.1 mm X 150 mm (Waters, Milford, MA, USA). The column was maintained at 45 °C and the flow-rate of 0.4 mL/min. The mobile phases consisted of (A) water with 10 mM ammonium formate, 0.125% formic acid, and (B) acetonitrile:water 90:10 (v/v) with 10 mM ammonium formate, 0.125% formic acid. The analytes separation was conducted following a stepwise gradient: 0-2 min 100% (B), 2-7.7 min 70% (B), 7.7-9.5 min 40% (B), 9.5-10.25 min 30% (B), 10.25-12.75 min 100% (B), 12.75-16.75 100% (B). A sample volume of 1 μL and 3 μL were injected for positive and negative mode, respectively. Negative and positive electrospray ionization (ESI) modes were applied with nitrogen serving as the desolvation gas and the collision gas. The parameters for detection of lipids were as follows: Curtain Gas: 35; CAD: High; Ion Spray Voltage: 4500 V; Source Temperature: 300°C; Gas 1: 60; Gas 2: 60; Declustering Potential: +/− 80 V, and collision energies +/− 10.

#### 3.2d) Metabolites: GC-MS conditions

A Leco Pegasus IV time of flight mass spectrometer coupled with Agilent 6890 GC equipped with a Gerstel automatic liner exchange system (ALEX) that included a multipurpose sample (MPS2) dual rail, and a Gerstel CIS cold injection system (Gerstel, Muehlheim, Germany) was used to complement HILIC metabolite analysis. The transfer line was maintained at 280 °C. Chromatography separation was achieved on a 30 m long, 0.25 mm i.d. Rtx-5Sil MS column (0.25 μm 95% dimethyl 5% diphenyl polysiloxane film) with the addition of a 10 m integrated guard column (Restek, Bellefonte PA) with helium (99.999%; Airgas, Radnor, PA, U.S.A.) at a constant flow of 1 mL/min. The oven temperature was held constant at 50°C for 1 min and then ramped at 20°C/min to 330°C at which it is held constant for 5 min. The GC temperature program was set as follows: 50°C to 275°C final temperature at a rate of 12 °C/s and hold for 3 minutes. The injection volume was 1 μL in splitless mode at 250 °C. Electron impact ionization at 70V was employed with an ion source temperature of 250°C. The scan mass ranged from 85 to 500 Da with acquisition rate of 17 spectra/second.

### 3.3) Statistical analysis

The statistical workflow used to find predictors of the CPET and HFBio tests, and reveal the metabolic pathways associated with HF, is depicted in **Fig 1**. Lipidomic and metabolomic data were presented as peak heights normalized by mTIC, a form of total ion chromatography with authentic metabolites that discounts abiotic artifacts. A compound is normalized by taking an individual peak height (mTICij) and dividing it by the product of average total peak heights and the sum of the specific compounds’ peak heights (mTICaverage x mTICsum,j). Using mTIC allows for the merger of the two databases. Outliers were eliminated using the Interquartile range (IQR) from a box-and-whisker plot with Turkey’s method detecting outliers as any value for the determined test outside of 1.5 times the IQR[21]. Metabolic data were filtered to exclude detected prescribed drugs, and from the lipidomic data only phospholipids, with detected fatty acids composition, were used in the analysis.

**Fig 1 –.**
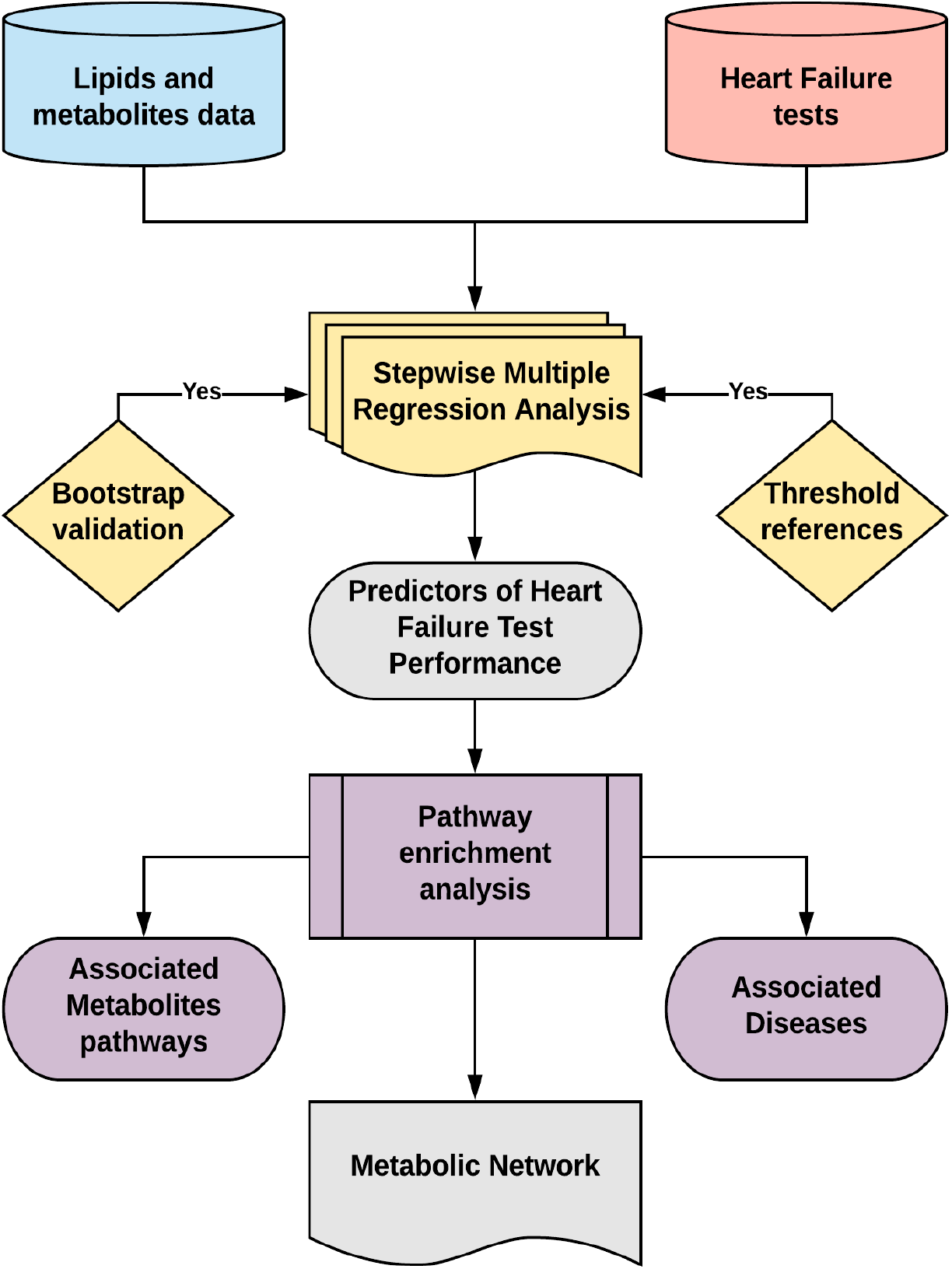
Statistical analysis workflow demonstrating the steps to find predictors of HF test performance using metabolomic and lipidomic data. Stepwise MRA was validated with bootstrap 95% confidence interval of the main regression estimates. Search of published threshold references confirmed the clinical importance of the models. Pathway enrichment analysis revealed the main metabolic pathways and associated diseases enriched using the set of metabolites predicting HF performance. A metabolic network was derived from the analysis confirming several metabolic dysfunctions related to HF described in the literature.

We utilized stepwise multiple regression analysis and Standard Least Squares methods to identify the set of metabolites which best associated with measures of cardiorespiratory fitness (peak oxygen consumption, oxygen uptake efficiency slope, exercise duration, VE/VCO2 slope) and established HF biomarkers of inflammation (CRP), fibrosis (galectin-3), and ventricular wall stress (N-terminal pro-BNP). Covariate bias was omitted from the MRA therefore eliminating risk of casual analysis and, because metabolic modulation was what was observed, unadjusted regression models were utilized in this study. Modulation was verified by comparison of these same measures with 316 lipids and 167 metabolites obtained from HF patients’ plasma.

MRA method was applied using the forward selection to enter the terms with the lowest p-values, allowing that at each step the candidate variable that increased the adjusted R^2^ the most was selected, until the model reached an adjusted R^2^ ≥ 0.8 and all the predictors had a statistically significant effect (p<0.05). All models reached a statistical significance of p<0.001. After the selection of predictors, the different models were standardized by mean centering to allow for the comparison of their coefficients, and the mean response of their dependent variables was estimated. The Root Mean Square Error (RMSE) was obtained and used as an estimate of each model’s fit. Coefficient of Variation (CV) was calculated as the ratio of RMSE to mean response of the dependent variable with its result suggesting good model fit and allowing for intermodal comparison.

To discover the metabolic pathways associated with HF, two methods of pathway enrichment analysis were performed. The first method was a Metabolic Pathway Analysis based on an over representation analysis with Fisher’s exact test algorithm to detect which metabolites were over-represented and significantly enriched. Coupled with this method, a pathway topological analysis with an out-degree centrality algorithm was used to measure the centrality of a metabolite in a metabolic pathway, estimating the mean importance of each matched metabolite impacting the pathway. The second method was a Metabolite Set Enrichment Analysis[22] that allows the incorporation into the analysis of pre-existing biological knowledge contained in metabolite-set libraries from studies of human metabolism. The analysis facilitated hypothesis generation and aided in interpretation of the metabolic models. Metabolite Set Enrichment Analysis used a reference metabolome from metabolite-set libraries to calculate a background distribution, and determine if the matched metabolite set in the model is more enriched for certain metabolites compared to random chance. The selection of pathway metabolites was based on the Kyoto Encyclopedia of Genes and Genomes (KEGG). To perform the study’s statistical analysis, multivariate linear regression was analyzed with JMP14Pro, and pathway enrichment analysis was performed with MetaboAnalyst 4.0.

## 4. Results

The study cohort included mostly African Americans with diabetes, hyperlipidemia, and hypertension (**Table 1**). The predictors of HF tests performance are listed in **Table 2**. The standardized regression coefficients suggest the expected elevation or decrease of lipids and metabolites in plasma based in their positive or negative values, although values higher than 0.3 were considered more reliable to indicate metabolic modulation.

**Table 1 –.** Demographics of the study cohort. Estimate’s data are presented as percentage or mean ± standard deviation.

| Covariates | Estimates |
| --- | --- |
| Age | 57.1 ± 11.7 |
| Male (%) | 75.5 |
| African American (%) | 79.6 |
| Coronary artery disease (%) | 36.7 |
| Diabetes (%) | 57.1 |
| Hyperlipidemia (%) | 63.3 |
| Hypertension (%) | 93.9 |
| Left ventricular ejection fraction (%) | 32.1 ± 9.5 |
| Left ventricular end-diastolic volume (mL) | 184.0 ± 62.5 |
| Left ventricular end-systolic volume (mL) | 127.0 ± 52.3 |
| Duke Activity Status Index | 28.4 ± 16.6 |
| Minnesota Living with Heart Failure Questionnaire | 58.6 ± 19.3 |
| Oxygen Uptake Efficiency Slope (L/min) | 1.8 ± 0.7 |
| Peak VO<sub>2</sub> (mL/min) | 14.0 ± 3.4 |
| VE/VCO<sub>2</sub> (mL/min) | 35.1 ± 7.3 |
| Exercise Duration (minutes) | 7.0 ± 2.6 |
| NT-proBNP (pg/mL) | 2650.3 ± 5475.1 |
| C-reactive protein (mg/L) | 8.2 ± 7.9 |
| Galectin-3 (pg/mL) | 21.4 ± 9.3 |

**Table 2 –.**
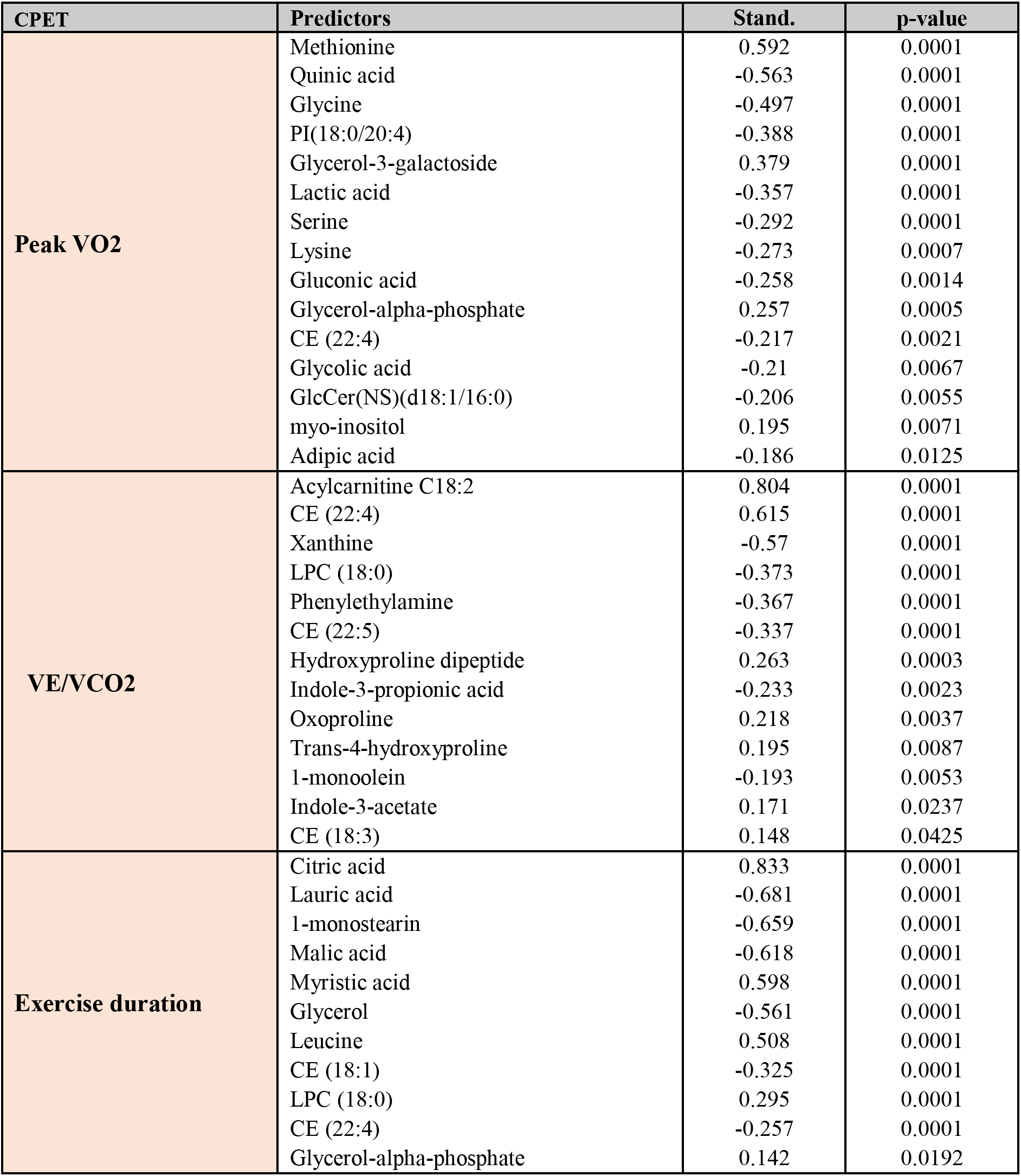
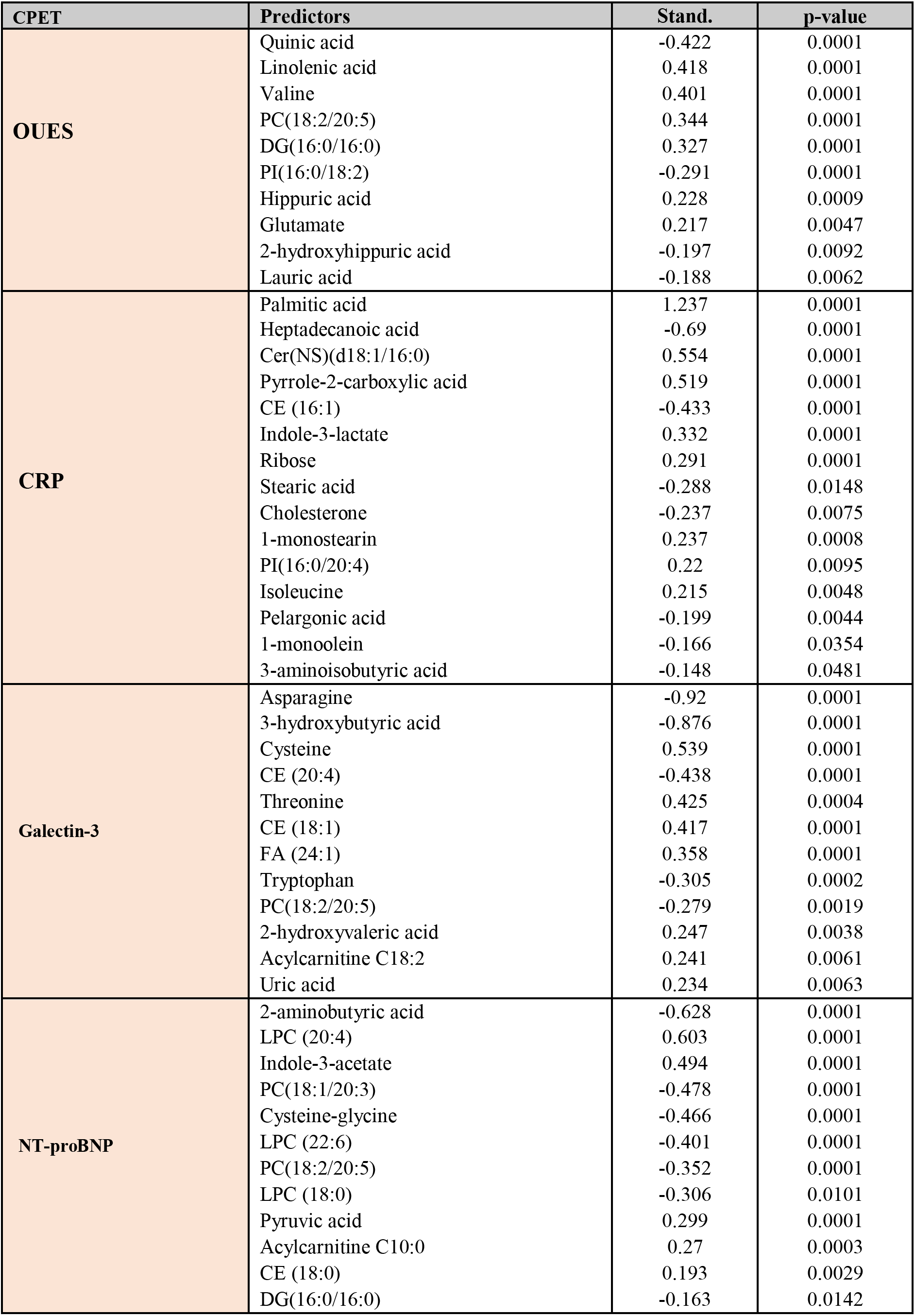
Predictors of cardiorespiratory fitness and traditional HF biomarkers. selected by multivariate linear regression. Standardized regression coefficient indicates the impact of one individual predictor over the specific test if all other predictors remain constant. Expected low values of exercise duration, OUES, and Peak VO2 and high values of VEVCO2, NT-proBNP, Galectin-3, and CRP predict poor test performance. Negative coefficient values are indicative of inverse correlation.

From the regression models it was possible to estimate the mean response of CPET and HFBio (**Table 3**) and compare them with threshold references published by other authors. The relative proximity of the tests mean response to these published threshold values indicates that metabolic modulation can indeed be assessed by HF tests performance. To be able to compare models we used the coefficient of variance, where the model with the smaller CV has predicted values that are closer to the actual values. VE/VCO2 and Peak VO2 (CV=0.07 and 0.08, respectively) showed the best metabolic predictive model for test performance. Galectin-3 (CV=0.13) also showed a reasonable predictive model, while CRP and NT-ProBNP (CV=0.31 and 0.29, respectively) were the least fit models.

**Table 3 –.**
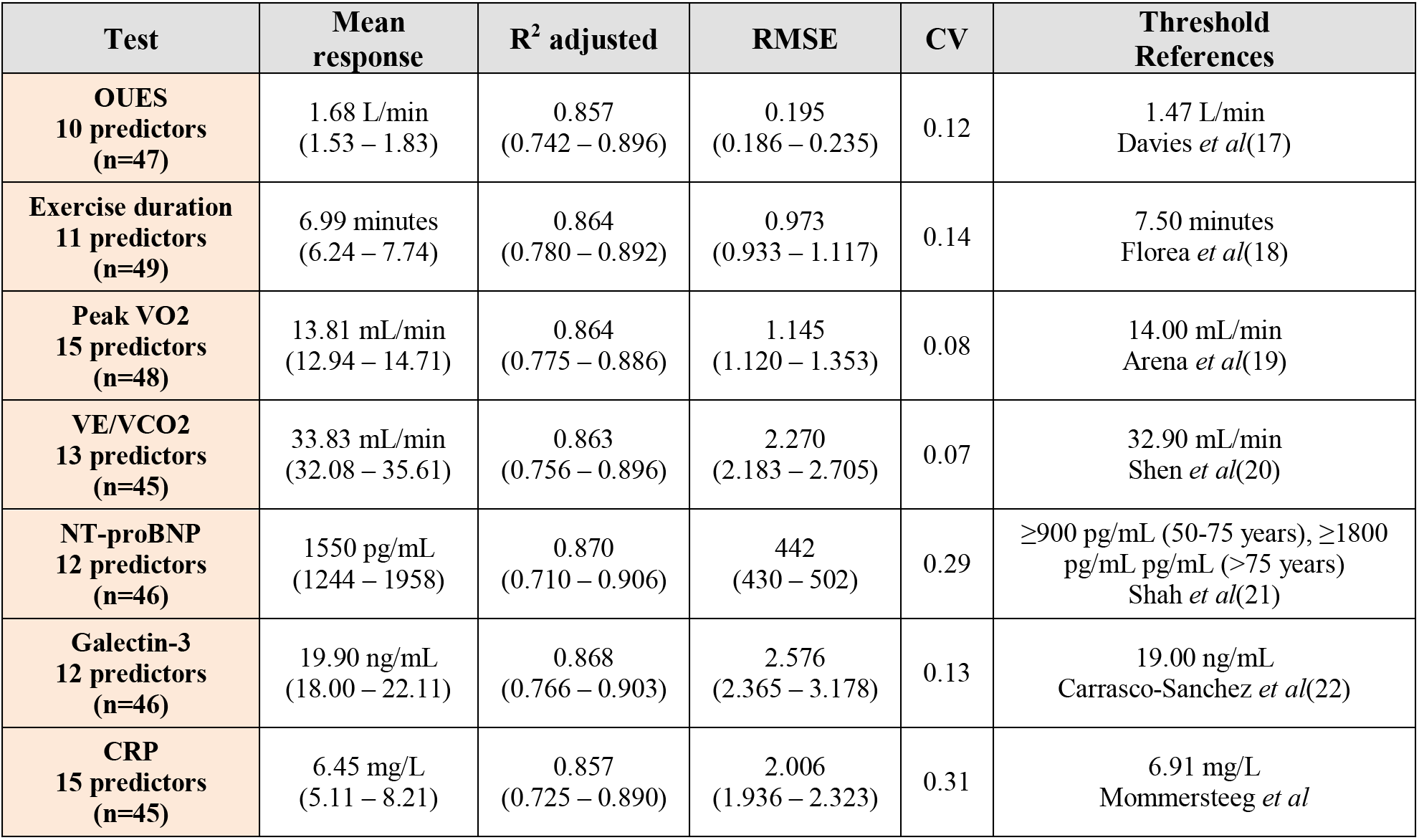
Estimates of cardiorespiratory fitness and HF biomarkers and comparison with literature threshold references. The mean response are values of the CPET and HFBio calculated from the regression parameters and a given value of the predictors that best fit the model. RMSE is presented as an absolute measure of fit in the same unit as the mean response. CV was calculated using the rate of RMSE by the mean response. Threshold references for test prediction of CHF outcome is presented as a measure of comparison of the mean response with peer reviewed publications. Bootstrap 95% CI are presented as a measure of validation, and are generally larger than 95% CI calculated from the actual data.

| Test | Mean response | R<sup>2</sup> adjusted | RMSE | CV | Threshold References |
| --- | --- | --- | --- | --- | --- |
| **OUES**<br>**10 predictors**<br>**(n=47)** | 1.68 L/min<br>(1.53 – 1.83) | 0.857<br>(0.742 – 0.896) | 0.195<br>(0.186 – 0.235) | 0.12 | 1.47 L/min<br>Davies *et al*(17) |
| **Exercise duration**<br>**11 predictors**<br>**(n=49)** | 6.99 minutes<br>(6.24 – 7.74) | 0.864<br>(0.780 – 0.892) | 0.973<br>(0.933 – 1.117) | 0.14 | 7.50 minutes<br>Floreas *et al*(18) |
| **Peak VO2**<br>**15 predictors**<br>**(n=48)** | 13.81 mL/min<br>(12.94 – 14.71) | 0.864<br>(0.775 – 0.886) | 1.145<br>(1.120 – 1.353) | 0.08 | 14.00 mL/min<br>Arena *et al*(19) |
| **VE/VCO2**<br>**13 predictors**<br>**(n=45)** | 33.83 mL/min<br>(32.08 – 35.61) | 0.863<br>(0.756 – 0.896) | 2.270<br>(2.183 – 2.705) | 0.07 | 32.90 mL/min<br>Shen *et al*(20) |
| **NT-proBNP**<br>**12 predictors**<br>**(n=46)** | 1550 pg/mL<br>(1244 – 1958) | 0.870<br>(0.710 – 0.906) | 442<br>(430 – 502) | 0.29 | ≥900 pg/mL (50-75 years), ≥1800<br>pg/mL (>75 years)<br>Shah *et al*(21) |
| **Galectin-3**<br>**12 predictors**<br>**(n=46)** | 19.90 ng/mL<br>(18.00 – 22.11) | 0.868<br>(0.766 – 0.903) | 2.576<br>(2.365 – 3.178) | 0.13 | 19.00 ng/mL<br>Carrasco-Sanchez *et al*(22) |
| **CRP**<br>**15 predictors**<br>**(n=45)** | 6.45 mg/L<br>(5.11 – 8.21) | 0.857<br>(0.725 – 0.890) | 2.006<br>(1.936 – 2.323) | 0.31 | 6.91 mg/L<br>Mommersteeg *et al* |

The diverse metabolites and lipids species compounding the CPET and HFBio prediction models are suggestive of the metabolic profile of the HF patient’s cohort. Therefore, they were used in the pathway enrichment analysis to indicate the more significant pathways impacting the prediction of HF patient’s performance. **Figure 2** shows that 13 pathways are significantly involved in prediction of HF test performance. Aminoacyl-tRNA and amino acid biosynthesis, amino acid metabolism, nitrogen metabolism, pantothenate and CoA biosynthesis, along with sphingolipid and glycerolipid metabolism, fatty acid biosynthesis, as well as glutathione metabolism and pentose phosphate pathway were revealed as the more relevant pathways.

**Fig 2 –.**
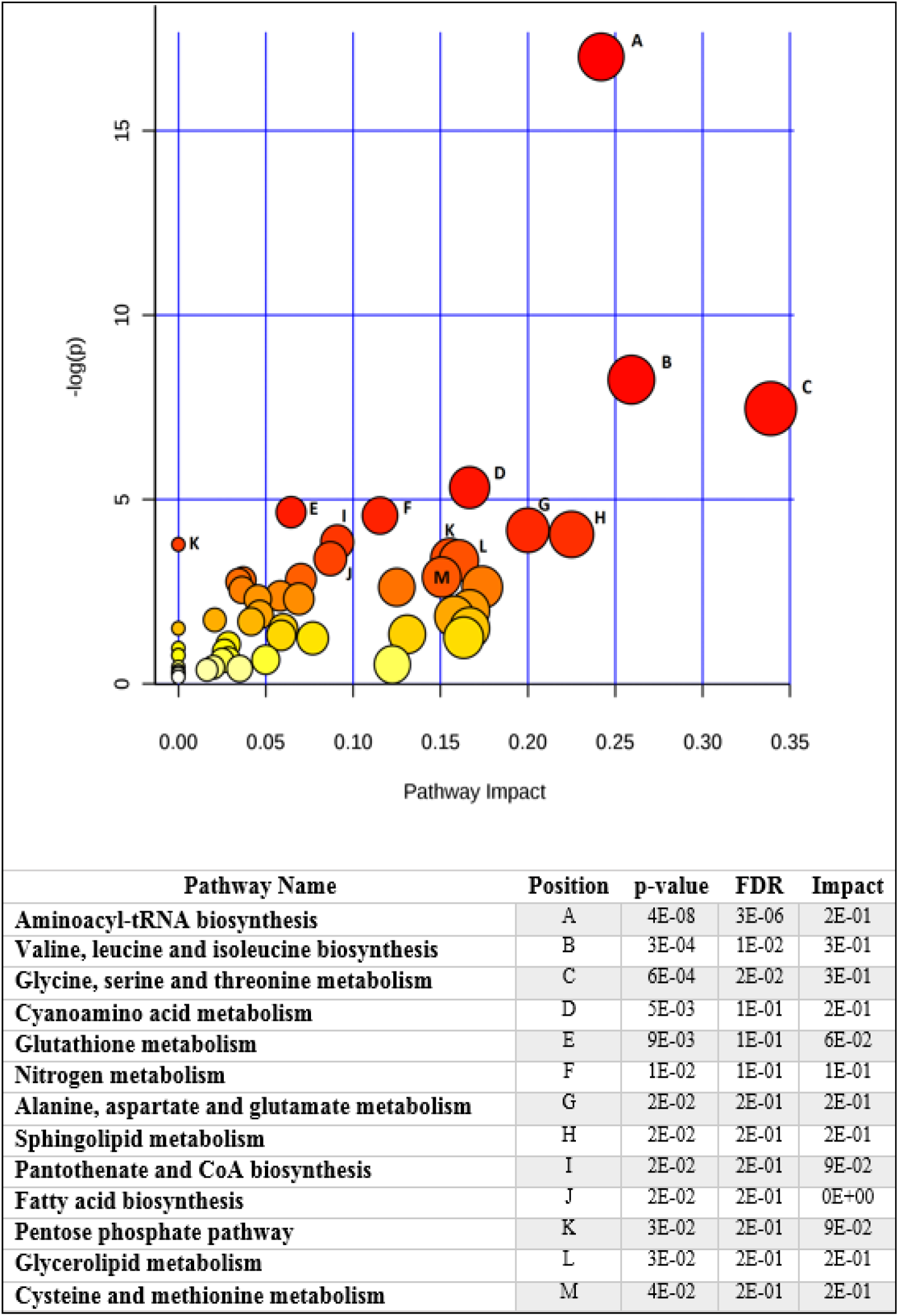
Metabolic pathway enrichment analysis shows the main pathways involved in heart failure test performance. The plot shows matched pathways according to the p-values from the pathway enrichment analysis and pathway impact values from pathway topology analysis. The pathways with the lowest p-values and highest match status (predictors present in the pathways) are listed in the table along with their FDR correction and impact score.

To explore the similarity of other diseases metabolic dysfunction to HF metabolic modulation, a pathway enrichment analysis was utilized (**Figure 3**). This analysis revealed 14 disorders statistically significant and with high impact that demonstrated similar metabolic perturbation to that observed by us for HF. Most of these identified diseases were related to brain dysfunction, such as acute seizure disorders and epilepsy-like metabolic profiles indicating the compromise of brain function. Enzymes deficiencies were also detected associated to lactic acidosis. Peritoneal dialysis and early markers of myocardial injury were present with lower impact. These results indicate that the metabolic profile of HF patients is mostly similar to brainlike dysfunction, and renal and cardiac abnormalities.

**Figure 3 –.**
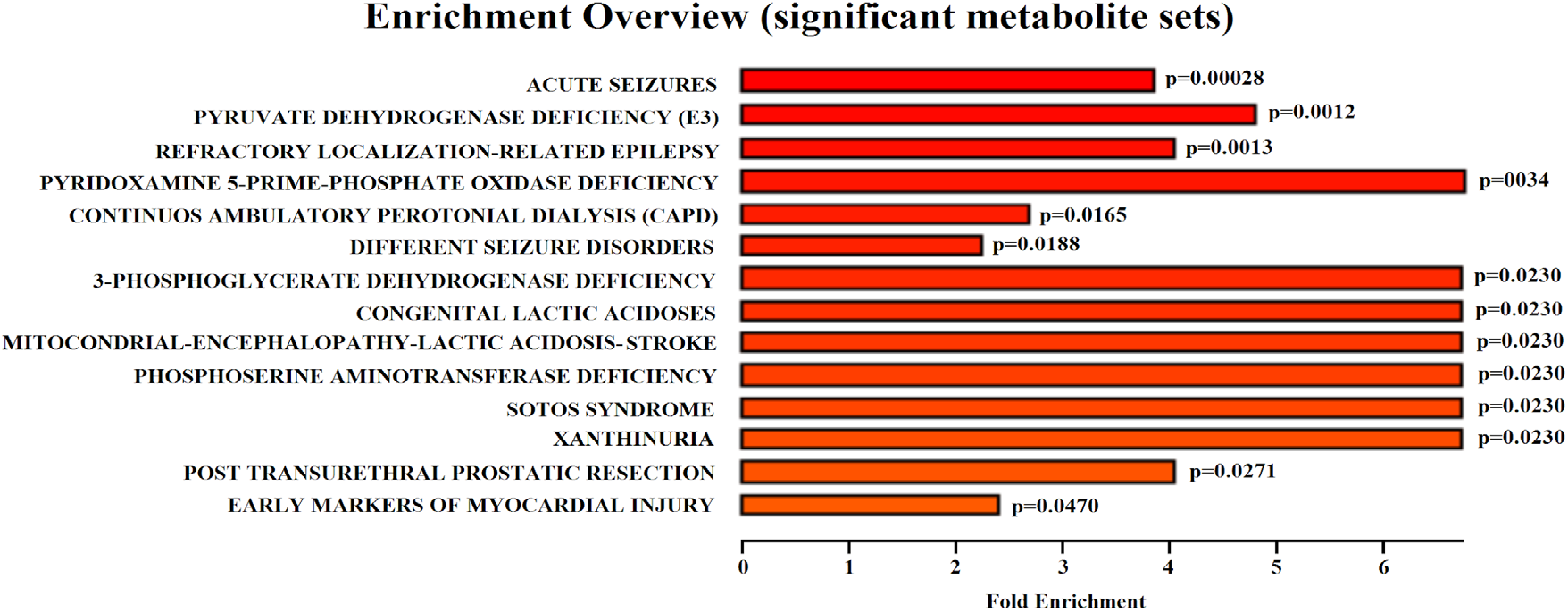
Diseases associated sets enrichment demonstrates the more important diseases presenting similar metabolic profile based in heart failure test performance. The majority of the statistically significant enriched diseases are related to brain dysfunction. Lactic acidosis-related diseases were also found with high impact in the analysis. One of the lowest impact diseases is early markers of myocardial injury.

Predictors of HF tests performance, supported by pathway enrichment analysis, were also used to propose a metabolic network suggestive of the metabolic modulations associated with HF test performance (**Figure 4**). In the proposed network, the positive or negative signs of the models’ coefficients were used as indication of metabolites elevation or decrease related to the expected poor test performance, respectively. The metabolic network was built linking relevant pathways revealed in the study, and the direction of pathways were suggested by metabolites elevation or decrease. Although the pathways are linked in the metabolic network, it is possible that not all metabolic pathways are occurring in the same tissues and organs.

**Figure 4 –.**
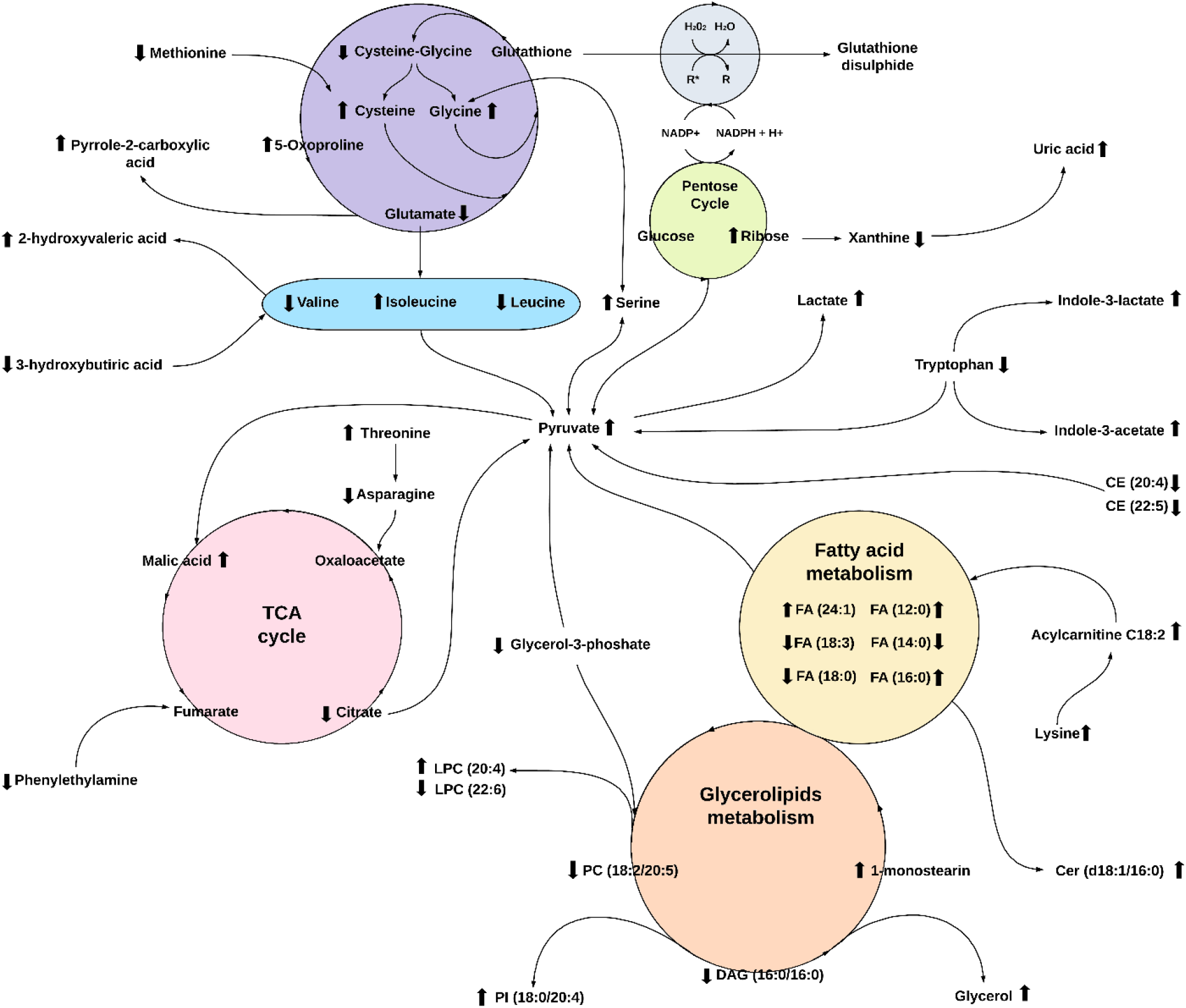
Intertwined metabolic network in Heart Failure. The metabolic changes affecting heart failure patients based in heart failure test performance includes glutathione anti-oxidative pathway, branched-chain amino acid (BCAA) biosynthesis, pentose cycle, tricarboxylic acid cycle (TCA), fatty acid (FA) metabolism, sphingolipids and glycerophospholipids metabolism, and tryptophan metabolism. Arrow represents predicted elevation or decrease variables in poor test performance. Only predictors with coefficients higher than 0.3 were used. Some metabolites not detected in the analysis were included in the figure to complement the metabolic pathways. Direction of pathways are proposed based in the metabolic modulation found in the study.

Considering the modulations in poor test performance, our results indicates an overall profile of oxidative stress, lactic acidosis, and metabolic syndrome, coupled with mitochondria dysfunction. There are signs of glutathione depletion, represented by decreased cysteine-glycine dipeptide and glutamate, and elevation of methionine, cysteine, glycine, and 5-oxoproline. The proposed oxidative stress could induce ribose elevation from the pentose phosphate pathway (PPP) and decreased xanthine and consequent elevation of uric acid in poor performance. Glutamate catabolism is indicated by elevation of downstream products such as pyrrole-2-carboxilic acid and hydroxyproline dipeptide.

Poor performance is associated with low levels of the branched-chain amino acids (BCAAs) leucine and valine, but elevation of isoleucine. Other amino acids modulations are elevated threonine and serine and decreased asparagine. Elevation of a compound similar to metabolites of BCAA catabolism, 2-hydroxyvaleric acid, was also detected. We also detected consumption of 3-hydroxybutyric acid, a ketone body linked to BCAAs metabolism, and decreased tryptophan levels. It appears that tryptophan catabolism is leading to elevation of indole-3-acetate and indole-3-lactate in an oxidatively stressed environment.

Pyruvate was elevated and appears to be a central metabolite linking several pathways. The decreased citrate in TCA cycle suggest that anaplerotic reactions are present. Therefore, the destiny of accumulated pyruvate could be responsible for malic acid elevation, but also responsible for elevation of lactate indicating lactic acidosis as a metabolic profile of poor HF test performance. Also in a link to fumarate in the TCA cycle, there was a decrease of phenylethanolamine.

Signs of fatty acids β-oxidation was inferred from the decreases in myristic acid (14:0), stearic acid (18:0), and linolenic acid (18:3). However, we found the levels of lauric acid (12:0), palmitic acid (16:0), and nervonic acid (24:1) were elevated in poor performance indicating special functions for these lipids. A sign that transport of fatty acids through the mitochondria membrane for β-oxidation is affected is the detected elevation of acylcarnitine C18:2 and lysine, a carnitine precursor, in the plasma. The analysis also suggests decreased levels of dietary polyunsaturated fatty acids corresponding to decreased levels of cholesteryl ester CE (20:4) and CE (22:5).

In the glycerolipids metabolism, elevation of lysophosphatidylcholine (LPC) carrying arachidonic acid (AA) and decrease of LPC carrying docosahexaenoic acid (DHA) suggests an imbalance toward accumulation of phospholipids containing pro-inflammatory fatty acids. This scenario is also supported by the detected decrease of phosphatidylcholine carrying eicosapentaenoic acid (EPA), PC (18:2/20:5), and elevation of phosphatidylinositol (PI) species containing omega-6 fatty acids represented by PI (16:0/18:2) and PI (18:0/20:4). We also observed decrease of DAG (16:0/16:0) that could be associated with the elevation of monoacylglycerol 1-monostearin and glycerol.

## 5. Discussion

Our analysis observed the presence of metabolic modulation dysfunction, characterized by indicators of oxidative stress, lactic acidosis, and amino acid and lipid metabolic disorders, within HF patients with poor cardiorespiratory fitness and unfavorable prognosis according to traditional HF biomarkers. It is known that patients with diabetes, hypertension, and hyperlipidemia presented a higher risk of oxidative stress and metabolic disorders due to decreased antioxidant defenses[23]. There is also a relationship between plasma total cysteine (tCys) and the risk of cardiovascular diseases and vascular toxicity of cysteine[24,25], as the fast auto-oxidation of cysteine can generate reactive oxygen species (ROS)[26]. High levels of cysteine and glycine are also associated with an elevated risk of developing metabolic syndrome[27]. A decreased methionine level is critical for anti-oxidative processes since methionine is a direct target of ROS, acting as a scavenger of free radicals[28,29]. Moreover, accumulation of 5-oxoproline in the blood could be responsible for metabolic acidosis associated with oxidative stress[30,31]. The reduction of glutamate can affect the brain’s activity, since glutamate acts as an excitatory neurotransmitter binding to the N-methyl-D-aspartate receptors and activate chloride ion channels[32].

As stated previously, it has been demonstrated that a reduction in the efficacy of amino acid metabolic activity with regards to poor test performance. Congestive HF (CHF) patients have reduced arterial amino acids that are related to HF severity[33]. Defective BCAAs catabolism and the elevation of branched-chain α-keto acids have also been associated with HF[34]. 2-hydroxyisovaleric acid has been reported in urine of patients with keto and lactic acidosis[35], as well as 3-hydroxybutyric acid has being found in patients with CHF[36]. Also, tryptophan degradation and elevation of indole-3-acetate suggest pro-inflammatory and pro-oxidant effects[37].

Pro-inflammatory and pro-oxidant effects were not the only physiological attributes we found. Our study also revealed that phenylethanolamine metabolism also was affected in poor performance. This metabolite is usually recognized as part of the phenylethanolamine N-methyltransferase (PNMT), an enzyme that converts norepinephrine to epinephrine. In a mouse model with knocked out PNMT, epinephrine-deficient’ mice had an exaggerated blood pressure response to exercise and reduced cardiac filling, indicating that epinephrine is required for maintaining normal cardiovascular function during stress[38].

Along with PNMT, elevated levels of fatty acid oxidation are a common metabolic disturbance in HF and other cardiopathies[39]. Activation of compensatory mechanisms such as PPP play a critical role in regulating cellular oxidative stress and lipids synthesis, although it can paradoxically feed superoxide-generating enzymes[40,41]. Ribose catabolism was linked to uric acid elevation in poor performance, and uric acid was associated with markers of metabolic syndrome and of systematic inflammation[42,43].

We detected a reduction of citrate and an elevation of malic acid in the TCA cycle, supporting the observation that the carboxylation of pyruvate to malate is an important anaplerotic reaction in HF[44]. In a study of hypertrophied rat hearts, the literature found that the rate of palmitic acid entering into oxidative metabolism was reduced by 23% compared to normal heart metabolic activity; the reduced rate of palmitate oxidation was balanced by a compensatory increase in anaplerotic flux, and the increased anaplerosis in hypertrophic hearts was fueled in part by increased carboxylation of the glycolytic pyruvate[45].

The elevated pyruvate in poor performance could be explained by the activation of fatty acid oxidation to produce acetyl-CoA, causing pyruvate oxidation inhibition in ischemic organs and driving pyruvate conversion to lactate[46]. Notwithstanding the detected oxidation of fatty acids, we found the levels of lauric acid (12:0), palmitic acid (16:1), and nervonic acid (24:1) were elevated in poor performance. Lauric acid elevation has being associated with an increased risk of coronary disease and ischemic stroke[47]. Palmitic acid is one of the more abundant fatty acids in human plasma and was the predictor with the highest impact for CRP. Wu *et al* reported that palmitic acid significantly stimulated in vitro CRP, TNF-α, and iNOS expression at the mRNA and protein levels in vascular smooth muscle cells[48]. Nervonic acid, a long-chain monounsaturated omega-9 fatty acid that undergoes β-oxidation in peroxisomes, plays an important role as an intermediate in the biosynthesis of nerve cell myelin[49]. Higher levels of nervonic acid have been positively associated with greater congestive HF and an increased risk of cardiovascular mortality, suggesting nervonic acid may pose as a cardiotoxin in humans[50]. Fatty acids like nervonic acid again have shown a significant role in HF modulation.

By the detection of elevated plasma acylcarnitine, we can infer that modulated fatty acids involved in HF β-oxidation are transported through the mitochondrial membrane. In studying the association of metabolites with adverse HF outcomes, Ahmad *et al* found that plasma acylcarnitine C18:2 was significantly higher in patients with end-stage HF[51]. The metabolic derangement in poor performance is also supported by the quantifiable differences in sphingolipid, glycerolipid, and glycerophospholipid metabolisms. In a rat model, myocardial Cer (d18:1/16:0) increased significantly 24 hours after acute myocardial infarction[52]. Elevation of pro-inflammatory lipids such as the specific LPC carrying arachidonic acid (AA) suggests an imbalanced accumulation of phospholipids containing omega-6 pro-inflammatory fatty acids[53]. This imbalance is further apparent by the presence of PC (18:2/20:5) and in our data with higher levels of phosphatidylinositol species containing omega-6 fatty acids PI(16:0/18:2) and PI(18:0/20:4). Phosphatidylinositol (PI) is especially abundant in brain tissue, along with a high AA content[54]. Biosynthesis of PI is linked to the reversible diacylglycerol (DAG) metabolism. DAG regulates the activity of protein kinase C that controls many key cellular functions, including ROS production[55]. Degradation of DAGs could be explained by the degradation of glycerolipids and glycerophospholipids from exacerbated oxidative stress[56].

## 6. Conclusion

Through our examination of metabolic dysfunction and HF patient test performances coupled to multiple regression research and analyses, we explored possible causal relationships among variables with respect to physiological context. In our study, MRA successfully demonstrated patterns of relationships that are consistent with interpretations reported by other authors, revealing the metabolic modulation associated with HF tests performance. Further studies using MRA with consideration to preexisting literature on physiology therefore could yield promising results, such as those found in this manuscript that could help improve our diagnoses of diseases and disorders.

## 7. Acknowledgement

Research reported in this publication was supported by research grants from National Institutes of Health under grant numbers HD087198 (to DSW), HL117026 (to BVT, AA) and from the Heart Failure Society of America (to LFB). The content is solely the responsibility of the authors and does not necessarily represent the official views of the National Institutes of Health. This work also received support via a Young Investigator Award from SCIEX for clinical lipidomic research (DSW).

## 8. Statement of Competing Financial Interests

All authors of this manuscript declare that they have no competing financial interests.

